# Effects of enriched biochar treatments on CO_2_ release, soil nitrate and ammonium, and wheat growth parameters in saline soils

**DOI:** 10.1101/2023.05.21.541629

**Authors:** Salahedin Moradi, Mirhassan Rasouli-Sadaghiani, Ebrahim Sepehr, Habib Khodaverdiloo, Mohsen Barin, Ramesh Raju Vetukuri

## Abstract

The effects of treatment with simple and enriched biochar on microbial respiration, nitrate and ammonium concentrations, and wheat growth parameters in saline soils were investigated using a completely randomized factorial experimental design with three replications, three soil salinity levels (1.5, 4.5 and 9 dS.m^-1^), and five biochar treatments including control, 2% simple (SB) or enriched biochar (EB) amendment, and 4% SB or EB amendment. The basal respiration rate and the concentrations of ammonium, and nitrate were measured at multiple time points. Additionally, total soil nitrogen, organic carbon, and microbial biomass carbon, microbial biomass nitrogen, and microbial biomass phosphorus were measured together with the height and fresh and dry weight of wheat after a 100-day growth period and at the end of the experiment. Salinity significantly affected basal respiration, nitrate and ammonium concentrations, plant height, and wet and dry weight. Biochar amendment significantly affected pH, basal respiration, nitrate and ammonium concentrations, total soil nitrogen, soil organic carbon, microbial biomass carbon, microbial biomass nitrogen, and microbial biomass phosphorus in both rhizosphere and non-rhizosphere soil, as well as wheat height, and wet and dry weight. The interaction between salinity and biochar significantly affected nitrate and ammonium concentrations and also plant height and fresh and dry weight. Finally, the effects of different biochar amendments and salinity levels on the basal respiration rate and the concentrations of nitrate and ammonium varied significantly over time. Overall, the results obtained show that biochar amendment can significantly moderate the adverse effects of soil salinity, especially if enriched biochar is used.

## Introduction

Saline soils are widely distributed in arid and semi-arid regions in over 100 countries around the world (Saifullah et al. 2018), including Iran. Salinity stress is an abiotic stress that affects many physiological parameters and processes in plants, including photosynthesis and levels of chlorophyll, proline, and carbohydrates. In addition, it causes osmotic changes around the roots and disrupts nutrient absorption (Nasraoui et al. 2013), reducing plant performance. Consequently, salinity is one of the most important factors reducing global agricultural product yields (Negrao et al. 2017). Salinity is also a major environmental stress for soil microorganisms that reduces the activity of fungi and bacteria, leading to reduced mineralization of carbon and organic nitrogen. Microorganisms in saline soils are adversely affected by the low osmotic potential of their surroundings, which causes their cells to lose water. Saline soils are also deficient in nutrients such as nitrogen and have high concentrations of sodium and chloride ions, leading to high ionic toxicity. These conditions result from the combined effects of multiple physical, chemical, and biological processes together with a lack of organic matter caused by poor plant growth.

The availability of nitrogen to plants may be very low under such conditions, which partly explains the low efficiency of nitrogen fertilizers in saline soils. Another contributing factor is that nitrification (Akhtar et al. 2012) and the mineralization of nitrogen (Irshad et al. 2005) and carbon (Setia et al. 2011) are slower in saline soils than in normal soils.

Increasing salinity reduces microbial biomass activity and thus reduces the rates of various biochemical processes. This reduces microbial respiration and carbon dioxide emissions, the mineralization of organic matter (Setia et al. 2011), and the rates of nitrification and ammonium production (Akhtar et al. 2012). Salinity also inhibits the decomposition of organic matter and thus slows down the nitrogen cycle, which can adversely affect plant nitrogen nutrition.

Biochar amendment can alter soil microbial activity and the rates of key processes in the nitrogen cycle (mineralization, nitrification, denitrification, and nitrogen fixation) because biochars can adsorb microbial signaling molecules and their pores provide protective habitats for microorganisms. Adding biochar to soil can thus increase rates of nitrification while also reducing emissions of N_2_O and the sublimation of ammonia, thereby increasing the bioavailability of nitrogen for plants and the rate at which native soil organic matter is mineralized. By retaining water in its pores, reducing soil density and increasing soil porosity, biochar causes the absorption of nitrifier inhibitors in the soil, as a result, it stimulates nitrification.

Because biochar amendment reduces nitrogen leaching, improves the biological properties of soil, and increases the activity of bacteria involved in the nitrogen cycle, it can increase both the storage and availability of nitrogen, thus potentially reducing the quantity of nitrogen fertilizer needed for crop growth. This work therefore investigated the interaction between biochar amendment and salinity to evaluate its effects on nitrogen and carbon mineralization, nitrification, and wheat growth and yield in saline soils.

## Material and methods

### Determination of soil characteristics

A greenhouse experiment with a factorial completely randomized design (CRD) was conducted to investigate how soil properties and wheat growth were affected by treating soil with simple and enriched biochar at loadings of 2% and 4% by mass. Soils of three differing salinity levels (1.5, 4.5, and 9 dS.m^-1^) were examined and the following variables were measured: CO_2_ emissions, ammonium and nitrate concentrations, microbial biomass carbon (MBC), microbial biomass nitrogen (MBN), microbial biomass phosphorus (MBP), and three wheat growth parameters - fresh and dry weight, and plant height. The soil’s physical and chemical properties were determined using standard methods (Table 1).

**Table 1.** Physical and chemical characteristics of the soil used in the experiment.

| Trait | Value |
| --- | --- |
| Sand (%) | 55 |
| Silt (%) | 26 |
| Clay (%) | 19 |
| BD (gr cm<sup>-3</sup>) | 1.3 |
| pH | 7.15 |
| ECe (dS m<sup>-1</sup>) | 0.58 |
| CCE (%) | 15 |
| OC (%) | 0.48 |
| N (%) | 0.035 |
| P (Olsen, mg kg<sup>-1</sup>) | 8.6 |
| K (mg kg<sup>-1</sup>) | 236 |
| Fe (DTPA, mg kg<sup>-1</sup>) | 7.23 |
| Zn (DTPA, mg kg<sup>-1</sup>) | 0.34 |
| Mn (DTPA, mg kg<sup>-1</sup>) | 6.83 |
| Cu (DTPA, mg kg<sup>-1</sup>) | 0.8 |

BD; Bulk Density, ECe; Electrical conductivity of saturated extracts, CCE; Calcium carbonate equivalent, OC; Organic carbon, N; Nitrogen, P; Phosphorus, K; Potassium, Fe, Iron, Zn; Zinc, Mn; Manganese, Cu; Copper

### Preparation of simple and enriched biochar

To prepare biochar, grape pruning residues were collected from vineyards. Enriched biochar was prepared by mixing simple biochar with 10% phosphoric acid in a 1:1 ratio and storing the mixture for 24 hours before mixing it with rock phosphate and cow manure in a ratio of 45:35:20 by mass. The mixed solids were then combined with distilled water in a ratio of 20:6 (water to mixture), after which the resulting suspension was placed in an oven and heated to 80°C for 24 hours followed by 220°C for one hour (Joseph et al. 2013; Chia et al. 2014). Selected physicochemical characteristics of the simple and enriched biochars are listed in Table 2.

**Table 2.** Selected physicochemical properties of the simple and enriched biochars.

| Trait | Value |  |
| --- | --- | --- |
|  | Simple biochar | Enriched biochar |
| C (%) | 76 | 68 |
| N (%) | 0.85 | 0.72 |
| O (%) | 15.8 | 16.6 |
| H (%) | 3.6 | 4.1 |
| P (g kg<sup>-1</sup>) | 1.6 | 4.8 |
| K (g kg<sup>-1</sup>) | 19.5 | 17.8 |
| Na (g kg<sup>-1</sup>) | 1.45 | 2.4 |
| Fe (mg kg<sup>-1</sup>) | 334 | 352 |
| Zn (mg kg<sup>-1</sup>) | 57 | 75 |
| Mn (mg kg<sup>-1</sup>) | 15.4 | 22.7 |
| Cu (mg kg<sup>-1</sup>) | 21.7 | 28.3 |
| pH | 8.2 | 5.1 |
| EC (dS m<sup>-1</sup>) | 2.41 | 3.6 |

C; carbon, N; Nitrogen, O; Oxygen, H; Hydrogen, P; Phosphorus, K; Potassium, Na; Sodium, Fe, Iron, Zn; Zinc, Mn; Manganese, Cu; Copper, EC; Electrical conductivity

### Establishing the desired salinity levels in the experimental soils

A mixture of salts chosen to mimic the ionic composition of the soils around Urmia Lake in Iran was used to establish the desired salinity levels in the experimental soils. The mixture consisted of MgSO_4_.7H_2_O, NaCl, Na_2_SO_4_ and CaCl_2_ in a ratio of 41.82:0.91:20.36: 36.91 by mass (Moradi et al. 2019). Pots with free drainage were used to prevent the accumulation of salts in the soil, and the soil salinity was measured weekly during the greenhouse experiment to detect deviation from the desired values, which were maintained by performing leaching as required.

### Plant cultivation and application of treatments

Air-dried soil samples were weighed and then poured into plastic bags. Based on preliminary concentration measurements and the critical nutrient levels determined for Iran by the Iranian Soil and Water Research Institute, four key nutrients (N, P, Zn, and Cu) were then added to the soil at concentrations of 60, 6.4, 0.66, and 0.2 mg kg^-1^, respectively, to prevent possible growth limitation. Biochar was then added to the relevant bags at the appropriate loadings and mixed uniformly with the soil, and the resulting mixtures were transferred to pots. Wheat seedlings were then placed in the pots and, to avoid sudden stress, salinity treatments were applied gradually after the plants were fully established. The pots were irrigated according to the FC. Soil respiration and the concentrations of ammonium and nitrate in the soil were measured at multiple points during the experiment using standard methods. At the end of the experiment, microbial biomass nitrogen (MBN) and microbial biomass carbon (MBC) were determined by the chloroform fumigation extraction method followed by back titration. Microbial biomass phosphorus (MBP) was calculated as the difference between the concentrations of extractable inorganic phosphorus in fumigated and non-fumigated soil (Brookes et al. 1982).

The gathered data were analyzed using the SAS software package (Statistical Analysis System, version 9.1 SAS Institute, Cary, NC, USA). Means were compared using Duncan’s multiple range test (DMRT) (P<0.05).

## Results

### Effect of salinity on basal respiration, soil concentrations of NH_4_^+^ and NO_3_^-^, and wheat growth parameters

Increased soil salinity was associated with reduced microbial respiration (Table 3). The basal respiration rate decreased gradually over time in all treatments.

**Table 3.** Effect of salinity (dS m^-1^) on basal respiration (mg CO_2_-C kg^-1^ Soil), NH_4_^+^ and NO_3_^-^ concentrations (mg kg^-1^), and selected wheat growth parameters

| Salinity | Basal Respiration | NH<sub>4</sub><sup>+</sup> | NO<sub>3</sub><sup>-</sup> | Height (cm) | FW (g) | DW (g) |
| --- | --- | --- | --- | --- | --- | --- |
| 1.5 | 50.6<sup>a</sup> | 77.2<sup>a</sup> | 6.6<sup>a</sup> | 40.6<sup>a</sup> | 5.1<sup>a</sup> | 1.1<sup>a*</sup> |
| 4.5 | 38.1<sup>b</sup> | 76.6<sup>a</sup> | 6.2<sup>a</sup> | 31.4<sup>b</sup> | 2.4<sup>b</sup> | 0.5<sup>b</sup> |
| 9 | 37.3<sup>b</sup> | 66.8<sup>b</sup> | 3.8<sup>b</sup> | 22.6<sup>c</sup> | 1.02<sup>c</sup> | 0.2<sup>c</sup> |

\*Values followed by the same letter in each column do not differ significantly at the P<0.05 level according to Duncan's test.
FW and DW indicates fresh and dry weight, respectively.

### Effect of biochar on basal respiration and concentration of NH_4_^+^ and NO_3_^-^ in soil

The biochar treatments increased the basal respiration rate (i.e., CO_2_ emissions from the soil). Throughout the experimental period, treatments with enriched biochar yielded higher basal respiration rates than those using simple biochar. Increasing the biochar loading from 2 to 4% also increased basal respiration for both types of biochar (Table 4).

**Table 4.**
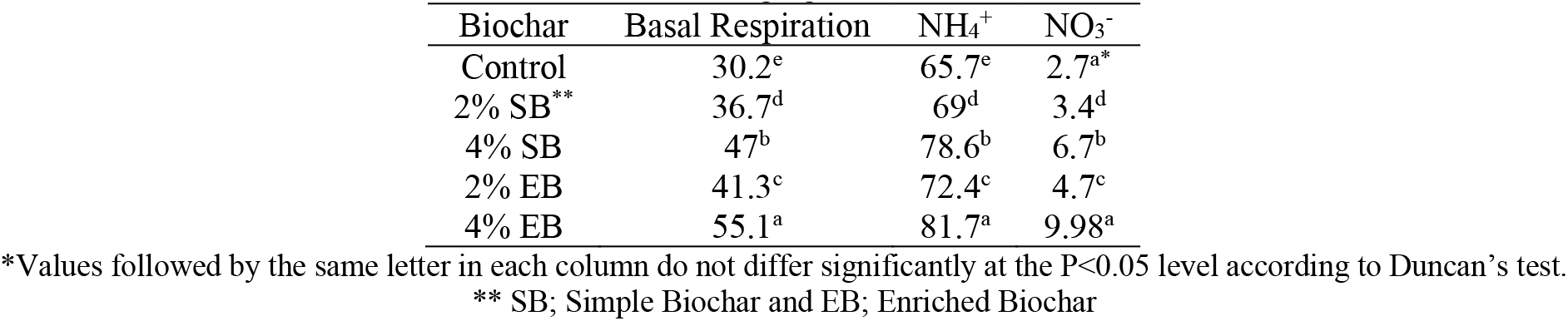
Effect of biochar loading (%) on basal respiration (mg CO_2_-C kg^-1^ Soil) and NH_4_^+^ and NO_3_^-^ concentrations (mg kg^-1^)

Treatment with simple and enriched biochar at the 2% and 4% levels increased the concentration of ammonium and nitrate in the soil, with higher concentrations generally being observed in soils amended at the 4% level.

### Effect of biochar on wheat growth parameters, soil pH, and concentrations of OC and N

Adding simple biochar increased the soil pH whereas enriched biochar reduced it (Table 5).

**Table 5.**
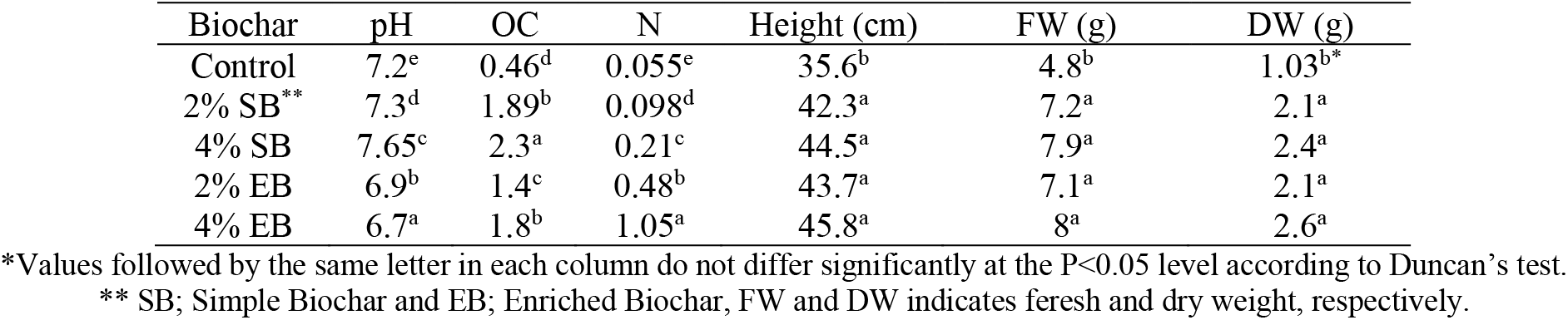
Effect of biochar (%) on soil OC and N concentration (%) and some wheat growth parameters

Biochar also increased the OC content and total N concentration of soils used to grow wheat. The OC content of soil treated with simple biochar was higher than that in control treatments and those using enriched biochar. Moreover, the OC content increased with the percentage of biochar in the soil; the OC content of soil amended with 4% simple biochar was 5 times that in the control treatment. Finally, the total N content of soil amended with enriched biochar exceeded that in simple biochar treatments and was up to 19 times higher than in control samples.

### Effect of biochar on MBC, MBN and MBP in rhizosphere and non-rhizosphere soil

The amounts of MBC, MBN, and MBP in both rhizosphere and non-rhizosphere soil were higher in the biochar treatments than in the control treatment (Table 6), indicating that biochar amendment enhanced microbial growth. Simple biochar had a stronger positive effect on MBC than enriched biochar, but the opposite was true for MBN and MBP. For both simple and enriched biochar treatments, the MBC, MBN, and MBP values observed in samples treated with 2% biochar did not differ significantly from those seen when the biochar loading was increased to 4%.

**Table 6.**
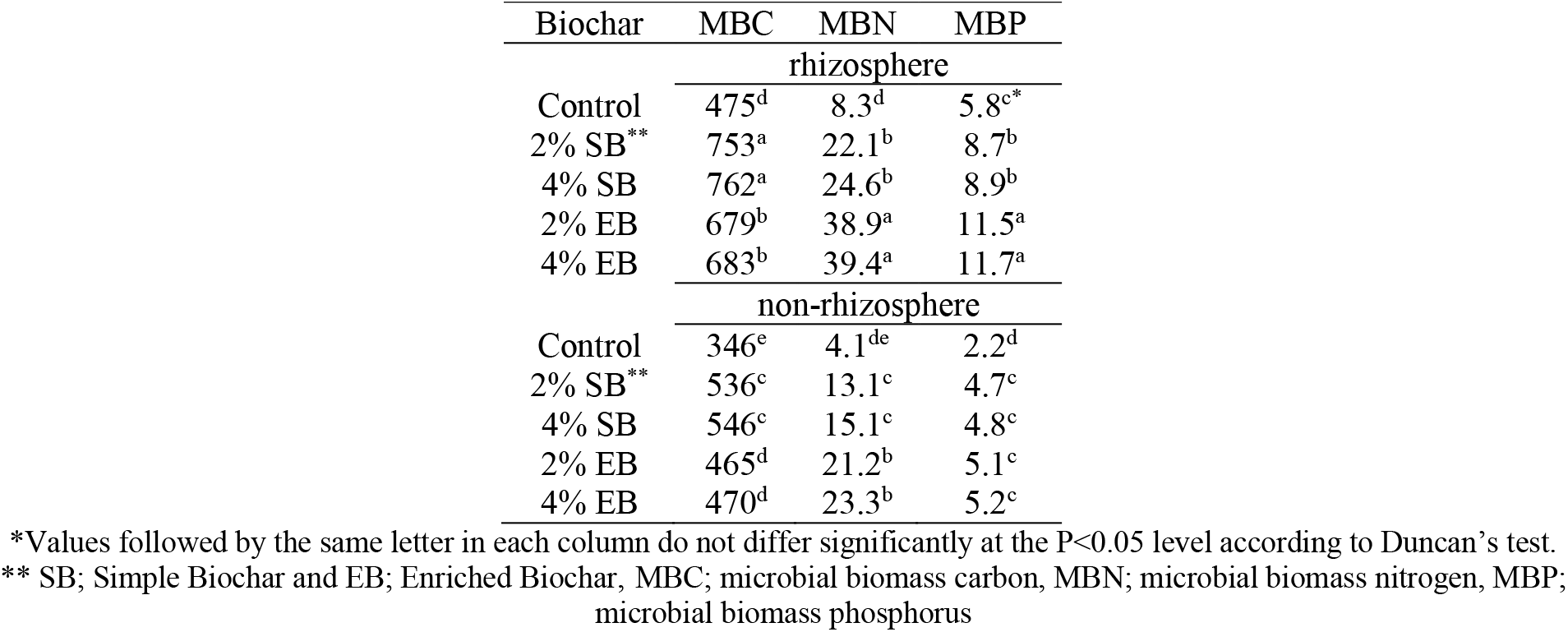
Effect of biochar loading (%) on soil MBC, MBN and MBP (mg.kg^-1^)

### Trends in basal respiration and soil concentrations of NH_4_^+^ and NO_3_^-^ over time

The basal respiration rate and the concentrations of NH_4_^+^ and NO_3_^-^ were highest on the 14^th^ day after the start of greenhouse cultivation (Fig. 1).

**Figure 1.**
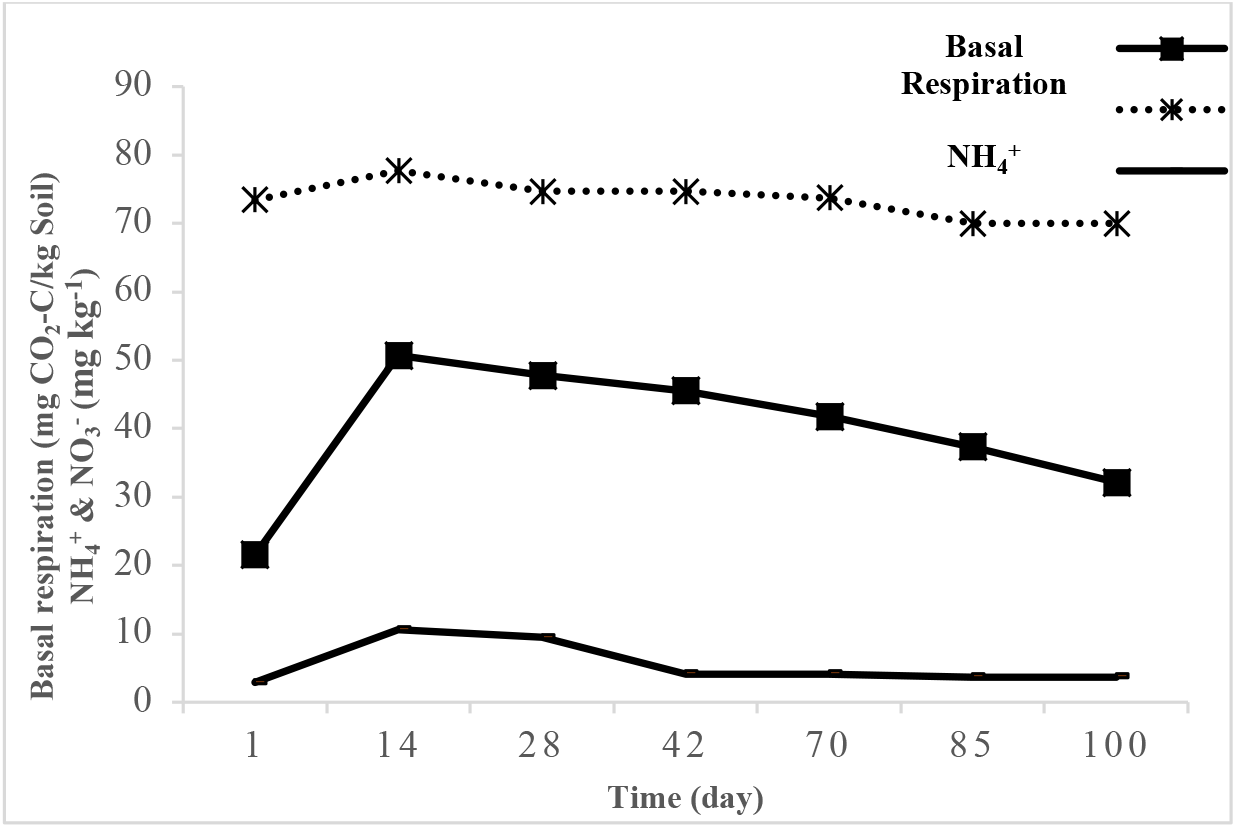
Changes in basal respiration and soil concentrations of ammonium (NH_4_^+^) and nitrate (NO_3_^-^) over time

### Interaction effects of salinity and biochar on NH_4_^+^ and NO_3_^-^ concentrations in soil and wheat growth parameters

Ammonium and nitrate concentrations decreased as soil salinity increased. At all salinity levels, biochar amendment increased the concentrations of NH_4_^+^ and NO_3_^-^ in the soil and enriched biochar had a stronger effect than simple biochar. The 4% biochar treatments generally had stronger effects than the 2% treatments but this difference was not always significant.

### Trends in basal respiration and concentrations of NH_4_^+^ and NO_3_^-^ over time in saline soil

The highest basal respiration rates and concentrations of NH_4_^+^ and NO_3_^-^ were observed in soils with a salinity of 1.5 dS m^-1^ on the 28^th^ day of the experiment; lower values were observed at later time points. However, the decrease in the NH_4_^+^ concentration started slightly later and only became apparent after the 42^nd^ day (Fig. 2), This may indicate that NH_4_^+^-producing microorganisms are more resistant to salinity than those producing NO_3_^-^.

**Figure 2.**
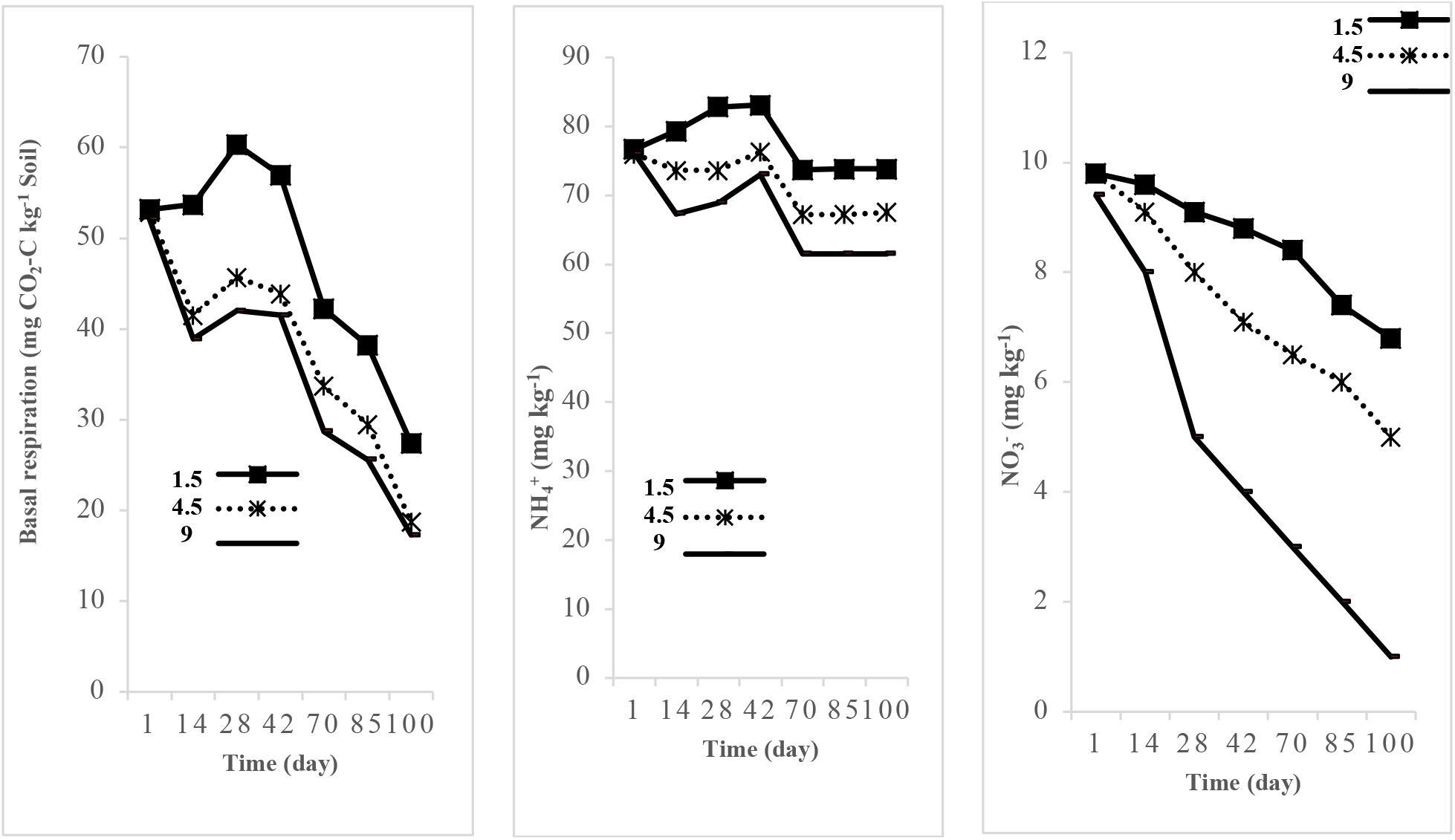
Trend in basal respiration and ammonium (NH_4_^+^) and nitrate (NO_3_^-^) concentrations in saline soils over time.

### Trends in basal respiration and NH_4_^+^ and NO_3_^-^ concentrations in biochar-amended soil over time

Biochar amendment generally increased the basal respiration rate and the concentrations of NH_4_^+^ and NO_3_^-^ relative to the control treatment. The highest values of all three variables were observed in the 2% biochar treatments on the 14^th^ day of the experiment. The lowest basal respiration rate was observed in the control treatment on the same day (Fig. 3).

**Figure 3.**
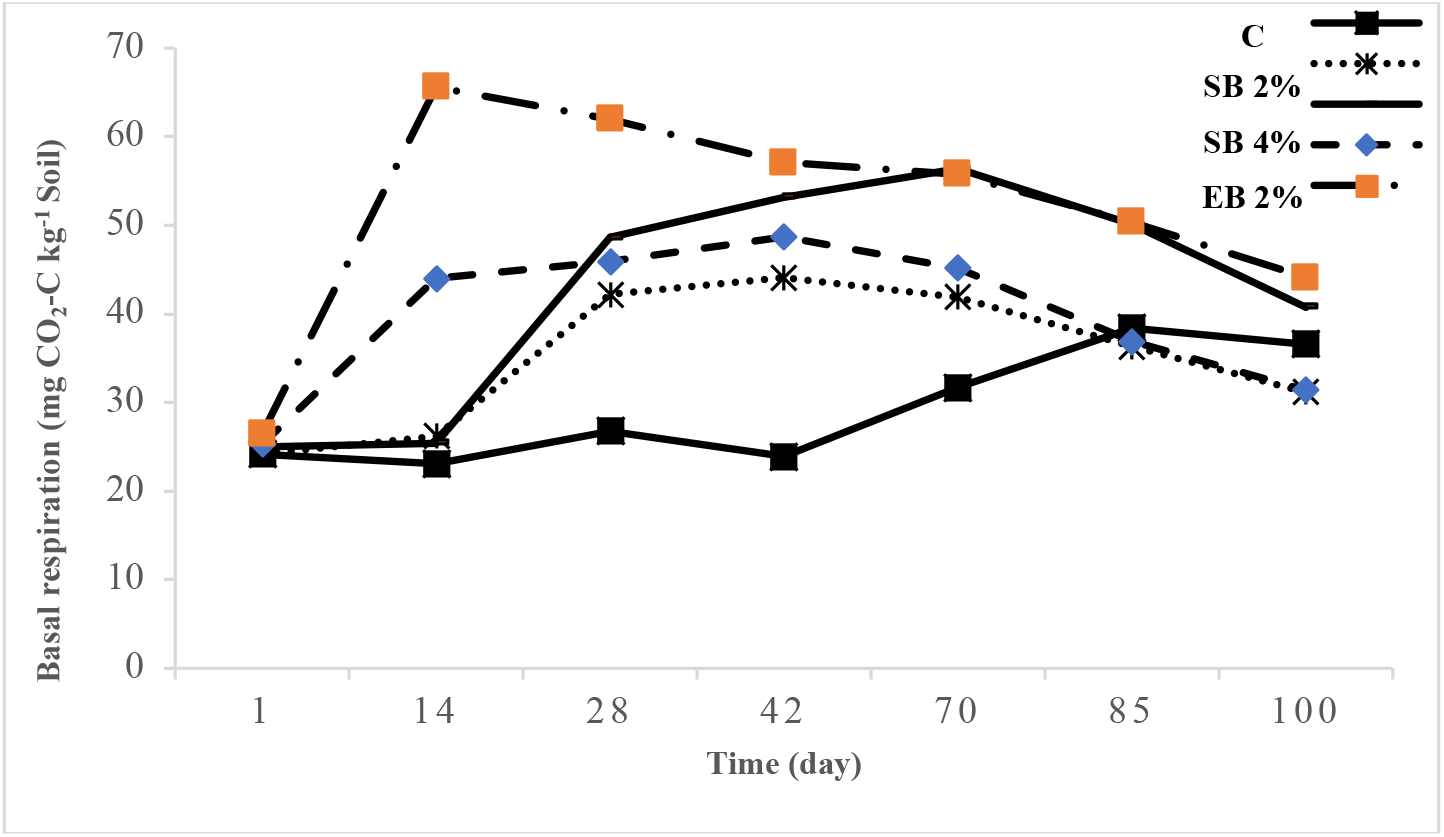
Trends in basal respiration in biochar-amended soil over time. SB; Simple Biochar. EB; Enriched Biochar.

The lowest NO_3_^-^ concentrations were observed in treatments without biochar 42 and 85 days after the start of the experiment. The lowest NH_4_^+^ concentration occurred in the control treatment on the 70^th^ day after the start of the experiment (Fig. 4).

**Fig4.**
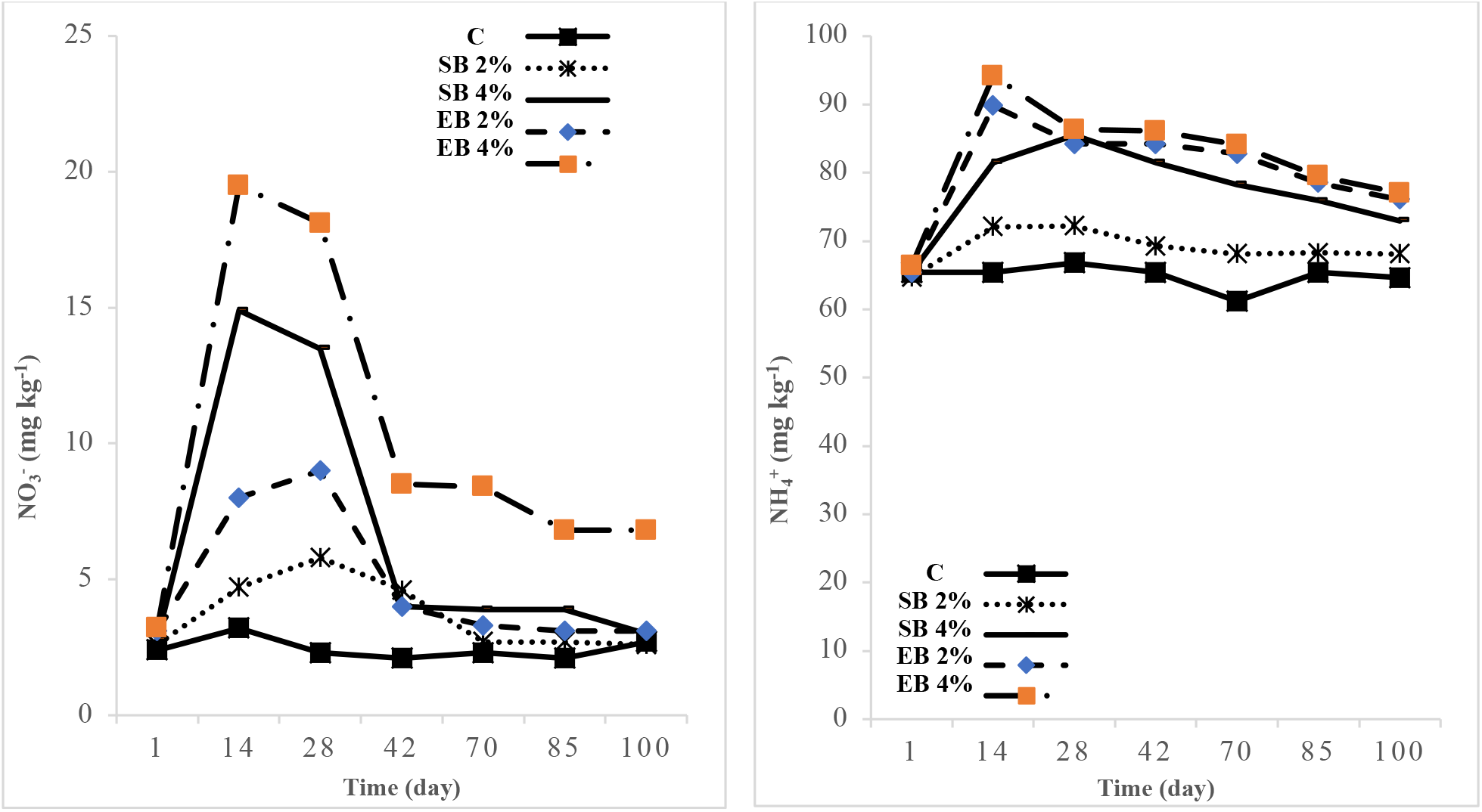
Trends in ammonium (NH_4_^+^) and nitrate (NO_3_^-^) concentrations in biochar-amended soil over time. SB; Simple Biochar. EB; Enriched Biochar

## Discussion

### Effects of salinity on basal respiration, soil concentrations of NH_4_^+^ and NO_3_^-^, and wheat growth parameters

Salinity is harmful to many microbial species and thus reduces the rate of microbial respiration. Accordingly, previous studies have found that increasing salinity reduces basal respiration under incubation conditions (Setia et al. 2011; Mousavi et al., 2023). Increased salinity also reduces the concentrations of ammonium and nitrate in soil under wheat cultivation; in this work, the concentrations of these ions were lowest in the most saline treatments (9 dS.m^-1^) and highest in the least saline ones (1.5 dS m^-1^). The rates of carbon and nitrogen mineralization and the degree of microbial activity thus all appear to be inversely correlated with soil salinity. This conclusion is supported by the findings of Akhtar et al. (2012), who reported that nitrification is strongly influenced by environmental stresses, especially salinity.

Our results also showed that salinity adversely affects various wheat growth indices, as can be seen in Table 3. For example, the fresh weight, dry weight, and height of wheat grown in the most saline soils examined in this work (9 dS m^-1^) were 23.89%, 81.81%, and 44.22% lower, respectively, than those for wheat grown in the least saline soils (1.5 dS m^-1^). Salinity thus appears to strongly affect the growth indices of aboveground wheat parts.

### Effects of biochar on basal respiration and soil concentrations of NH_4_^+^ and NO_3_^-^

Biochar amendment increased the concentrations of ammonium and nitrate in the soil when compared to control treatments, and this effect was more pronounced for enriched biochar than for simple biochar. The strong effect of enriched biochar is presumably due to its high total nitrogen content. Adding biochar to soil also increases microbial respiration and stimulates microbial activity in soil because the surface of biochar is rich in labile carbon and nutrients. However, these nutrients are consumed over time, so the positive effect of biochar amendment on microbial respiration diminishes over time (El-Naggar et al. 2015).

Enriched biochar has a particularly high nutrient content, which may partly explain the high release of CO_2_ observed in enriched biochar treatments. In keeping with this hypothesis, Sarkhot et al. (2012) reported that enriching biochar with manure increased its nutrient content and that amendment with such enriched biochar effectively increased soil microbial activity. Because enriched biochar also contains essential nutrients such as phosphorus and nitrogen, it can stimulate various microbial processes in the soil – for example, Nelissen et al. (2012) observed a short-term stimulation of nitrification following biochar amendment.

### Effects of biochar on pH, soil OC and N contents, and wheat growth parameters

Biochar is relatively rich in alkaline elements, which may be why soil pH has been reported to increase following treatment with simple biochar (Kookana et al. 2011). However, the effects of biochar on soil pH depend strongly on the material from which the biochar is prepared and the pyrolysis conditions (Yadav et al. 2016). For example, Streubel et al. (2011) reported soil pH increases of 0.1-0.8 units following treatment with different biochars. In contrast, amendment with enriched biochar may reduce soil pH because phosphoric acid is used in the enrichment process: Sun et al. (2016) reported a soil pH reduction of 0.2 units following the addition of enriched straw and wheat stubble biochar.

Under certain soil conditions, some of the C and N in added biochar may be released, increasing the concentrations of these nutrients in the soil. Enriched biochar has a particularly strong positive effect on soil nitrogen, possibly because the nutrients in simple biochar tend to be fixed whereas those in enriched biochar are more readily available to microorganisms. This effect seems to be especially pronounced for biochar enriched with manure. The presence of wheat may also affect certain soil characteristics. For example, plant roots can affect nutrient concentrations by releasing compounds such as organic acids into the soil.

Biochar is an organic material with a high carbon content, so it is unsurprising that its addition to soil tends to increase soil OC levels, as noted by Wu et al. (2016). Additionally, Zheng et al. (2013) have reported increased soil N levels following biochar addition, and Clough et al. (2013) argued that this increase can be attributed to biochar’s high specific surface area and CEC, both of which disfavor denitrification.

In our experiments, all of the tested biochar treatments improved the measured wheat growth indices, and the results presented in Table 5 show that there were no statistically significant differences between the improvements achieved under the different biochar amendment regimes. This is consistent with the findings of Saifullah et al. (2018), who stated that biochar has many beneficial effects that increase crop yields, one of which is a tendency to increase nutrient availability in the soil. Similar conclusions were reported by Zhao et al. (2014).

### Effects of biochar on MBC, MBN, and MBP in rhizosphere and non-rhizosphere soil

The MBC, MBN, and MBP in the rhizosphere exceeded those in non-rhizosphere soil. Increases in microbial biomass following biochar amendment have been attributed to its content of labile carbon (Luo Gu 2016). Additionally, the accumulation of biochar on the soil surface provides a substrate for microorganisms and thus increases the carbon content of the microbial biomass.

### Trends in basal respiration and soil concentrations of NH_4_^+^ and NO_3_^-^ over time

The basal respiration rate and the concentrations of nitrate and ammonium both tended to decrease over time in our experiments. Yao et al. (2010) found that C and N may become immobilized over time, reducing the availability of their mineral forms in the soil. Moreover, Sun et al. (2016) showed that the mineralization of organic compounds in the soil may initially increase and then decline following biochar addition rather than decreasing linearly. These findings are consistent with our results.

### Combined effects of salinity and biochar amendment on soil NH_4_^+^ and NO_3_^-^ concentrations and wheat growth parameters

Biochar amendment can help balance ionic ratios and thereby enhance rates of microbial mineralization processes that increase soil concentrations of NH_4_^+^ and NO_3_^-^. For example, Saifullah et al. (2018) found that adding biochar to saline soils increased their fertility and the accessibility of nutrients (especially N) to soil microbes, leading to activation of the sensitive soil microbial population and increased C and N mineralization. Similarly, Zhao et al. (2014) reported that the organic matter in biochar can increase nitrification by increasing the availability of key nutrients in the soil, and Ball et al. (2010) found that biochar formed during forest fires increases microbial activity and nitrification in soils.

Our results revealed that the interaction between biochar and salinity significantly affected the height and fresh and dry weight of wheat. As shown in Table 7, the values of all these wheat growth parameters were lowest in the biochar-free control treatment with a salinity of 9 dS m^-1^ and highest in the 4% enriched biochar treatment with a salinity of 1.5 dS m^-1^; the differences between these two treatments with respect to fresh weight, dry weight and wheat height were 86.9%, 92.4%, and 49.8%, respectively. Pandit et al. (2018) reported that biochar increased the yield and growth of corn by increasing the soil’s moisture retention capacity and moderating the effects of environmental stress, while Wong et al. (2009) stated that biochar treatment increases plant growth and yields by increasing soil moisture and the accessibility of micronutrients.

**Table 7.**
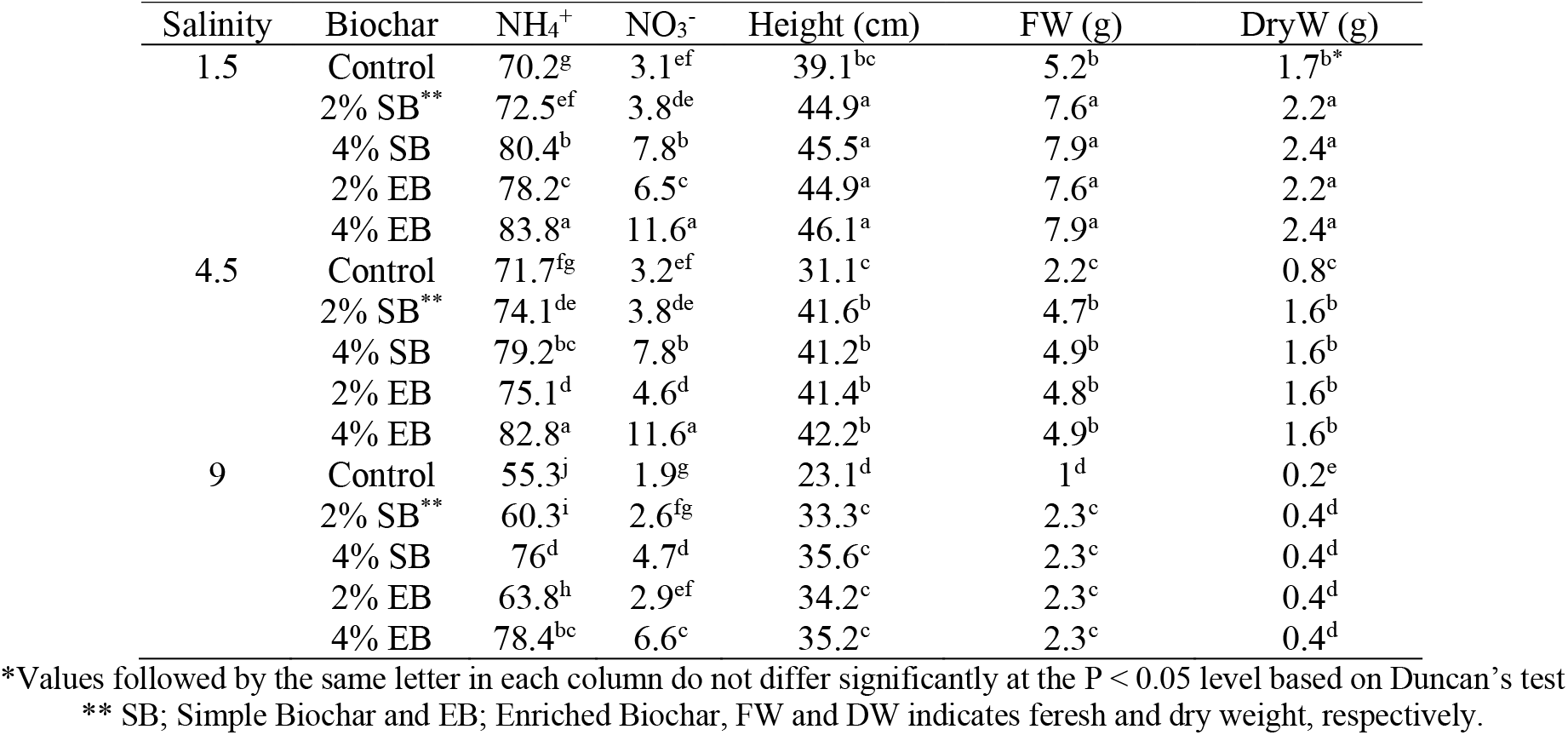
Effect of salinity (dS.m^-1^) and biochar (%) on NH_4_^+^ and NO_3_^-^ concentrations and wheat growth parameters

### Trends in basal respiration and concentrations of NH_4_^+^ and NO_3_^-^ in saline soil over time

The severity of the damage caused to soil microorganisms by salinity increases over time, leading to gradually decreasing rates of microbial respiration and production of NH_4_^+^ and NO_3_^-^ (Akhtar et al., 2012). Akhtar et al. (2015) also showed that the magnitude of the decline in NO_3_^-^ production is greater than that seen for NH_4_^+^ production, which they attributed to changes in the mineralization processes of organic matter. Further research on soil microbial communities will be needed to clarify these changes.

### Trends in basal respiration and concentrations of NH_4_^+^ and NO_3_^-^ over time in biochar-amended soil

The use of organic amendments such as biochar can be useful in situations where nitrification in agricultural soils has been reduced or halted by environmental stresses such as salinity or drought because biochar addition changes the properties and conditions of soil in ways that can increase the mineralization of organic matter (Wang et al. 2018). In particular, biochar can change the chemical forms of N in soil by increasing N fixation and NH_4_^+^ storage (Zheng et al. 2013) while reducing N_2_O release (Xu et al. 2014) and ammonia sublimation (Jones et al. 2012), leading to improved N availability for plants (Zheng et al. 2013).

Biochar is a very stable material with high resistance to decomposition (Awad et al. 2013). A major reason for its positive effect on nitrogen mineralization in the short term is that it can weaken the effects of nitrification inhibitors such as phenols (DeLuca et al. 2006). However, in certain cases it can also reduce the rate of carbon mineralization (Zimmerman et al., 2011). Lehmann et al. (2011) attributed this primarily to the adsorption of labile carbon to biochar surface and within its pore network. In another example, Maucieri et al. (2017) found that salinity and biochar amendment reduced microbial CO_2_ production by 10% and 24%, respectively, relative to controls. In another case, the mineralization of organic carbon in saline soils increased substantially following the addition of biochar but then decreased gradually over time (Sun et al. 2016). This probably occurred largely because biochar contains relatively little labile carbon (Major et al. 2010; Jones et al. 2011); once this food source is consumed, labile carbon becomes a limiting factor on microbial activity, leading to a decline in basal respiration (Kuzyakov et al. 2014).

## Conclusion

Salinity indirectly affects the nitrogen status of soils and reduces their content of available nutrients. Biochar amendment can alleviate these problems because biochar creates favorable conditions for nitrifying bacteria by altering the soil pH and adsorbing nitrification-inhibiting organic compounds, leading to short-term increases in the rate of nitrogen mineralization. Biochar can also be an easily accessible source of carbon that increases carbon and nitrogen mineralization in soil by stimulating microbial activity. However, over longer periods added biochar may reduce the concentrations of nitrate and ammonium in soils because it has a high specific surface area and a high ion exchange capacity. Under saline conditions, biochar can weaken osmotic effects and specific ionic toxicity by adsorbing salts and balancing ionic ratios, thereby creating conditions that allow soil microorganisms to be more active. Biochar enriched with other organic and inorganic compounds seems to be particularly effective at increasing microbial activity because it has a greater quantity and diversity of nutrients than simple biochar.

A common problem when applying biochar to non-acidic soils is that it may increase their pH. However, amendment with enriched biochar tends to reduce soil pH, making it an attractive option for improving the condition of such soils, especially since pH reductions may increase the accessibility of nutrients including phosphorus and micronutrients. In addition, amendment with enriched biochar has been linked to reductions in sodium concentrations. The results presented here show that enriched biochar amendment can effectively modulate the adverse effects of salinity in alkaline soils. This is important in the Iranian context because most Iranian soils are alkaline, which limits the usefulness of amendment with simple biochar. Enriched biochar treatment could thus be an attractive option for increasing the fertility of Iran’s soils while also reducing their pH.

## Data Availability Statement

The data presented in this study are available on request from the corresponding author.

## Acknowledgments

RRV acknowledges support from FORMAS (2019-01316) and the Swedish Research Council (2019–04270), Novo Nordisk Fonden (0074727), Carl Tryggers Stiftelse (CTS 20:464) and SLU’s Centre for Biological Control.

## Author Contributions

conceptualization: M.H.R.-S, E.S., H.K and M.B.; formal analysis: S.M.; utilization of software: S.M.; methodology: M.H.R-S., E.S., H.K and M.B.; investigation: S.M.; validation: M.H.R-S., E.S. and M.B.; data curation: M.H.R. original draft preparation: S.M.; review and editing: M.H.R-S., H.K., M.B. and R.R.V.. All authors have read and agreed to the published version of the manuscript.

